# Anesthesia alters cortical spike sequences in rodent visual cortex

**DOI:** 10.1101/2022.12.13.520271

**Authors:** Sean Tanabe, Heonsoo Lee, Shiyong Wang, Anthony G. Hudetz

## Abstract

Recurring spike sequences are thought to underlie cortical computations and may be essential for information processing in the conscious state. How anesthesia at graded levels may influence spontaneous and stimulus-related spike sequences in visual cortex has not been systematically examined. We recorded extracellular single-unit activity in the rat primary visual cortex *in vivo* during wakefulness and three levels of anesthesia produced by desflurane. The latencies of spike sequences within 0~200ms from the onset of spontaneous UP states and visual flash-evoked responses were compared. During wakefulness, spike latency patterns linked to the local field potential theta cycle were similar to stimulus-evoked patterns. Under anesthesia, spontaneous UP state sequences differed from flash-evoked sequences due to the recruitment of low-firing excitatory neurons to the UP state. Flash-evoked spike sequences showed higher reliability and longer latency when stimuli were applied during DOWN states compared to UP states. At deeper levels, anesthesia altered both UP state and flash-evoked spike sequences by selectively suppressing inhibitory neuron firing. The results reveal anesthesia-induced complex changes in cortical firing sequences that may influence visual information processing.

## INTRODUCTION

Recurring spike sequences are thought to underlie cortical computations and may be essential for information processing, predictive computations, memory consolidation, navigation, and replay (Luczak et al., 2015; Maboudi et al., 2018; Capone et al., 2019; Mackevicius et al., 2019; Luczak et al., 2022). Upon the presentation of a sensory stimulus, a set of cortical neurons respond with an increase of their firing rate with diverse latencies (Luczak et al., 2007). Such latencies form a reproducible sequence that is apparently conserved across stimulus trials, stimulus properties and cortical states, suggesting that they are determined by intracortical circuits (Watts and Thomson, 2005; Luczak et al., 2007; Luczak et al., 2009). The latency and firing pattern of neurons within the first 100ms post-stimulus period are similar between simple stimuli such as auditory tones and natural sounds (Luczak et al., 2009), whereas the later component of firing response is more diverse and thought to code complex stimulus properties (Luczak et al., 2015).

Similar to the stimulus-induced response, a sequential onset of firing occurs when neurons fire in spontaneous bursts (Luczak et al., 2007). This is generally observed under anesthesia at a level that facilitates intense, transient firing of neurons with a well-defined onset from a silent baseline. UP states in anesthesia are different from the those during wakefulness; in the latter, spontaneous activity is higher (Vizuete et al., 2014), and population spike sequences are difficult to discern due to their frequent overlap (Luczak et al., 2013).

The administration of anesthetics provides a well-controlled pharmacological approach to alter the state of cortex, neuronal excitability, population activity, and sensory function (Vizuete et al., 2012; Vizuete et al., 2014; Lee et al., 2020b; Lee et al., 2020a). Anesthetic agents inevitably modify the neuronal circuitry to slow electrophysiological oscillations (Purdon et al., 2015) and decrease the excitatory and inhibitory spike rates (Taub et al., 2013). Most anesthetic drugs exert their hypnotic effect via potentiating inhibitory transmission at GABAA receptors (Alkire et al., 2008), potentially altering the excitatory-inhibitory balance (Taub et al., 2013) and modifying the spike sequences transitioning from DOWN to UP states (Hasenstaub et al., 2007). Prior studies of spike latency sequences involving anesthesia mostly used urethane that has mixed pharmacology or a few other anesthetic combinations that suggested some agent-dependent effects (Luczak et al., 2007). For example, putative pyramidal neurons and interneurons responded to auditory stimuli with similar latency under urethane anesthesia (Luczak et al., 2009), but not under ketamine-xylazine anesthesia, when interneuron latency was approximately 20% shorter than pyramidal neuron latency. Whether such a difference exists with other types of anesthetics, including the pharmacologically different class of inhalational agents that preferentially modulate GABAA receptors, has not been evaluated. Thus, there is a need to extend the existing findings with using an inhalational anesthetic used to suppress consciousness in the clinical setting. Also, because many of the synaptic effects of anesthetics are dose-dependent, a better understanding of their influence on spike sequences necessitates an investigation of dose-dependent changes during spontaneous activity and sensory stimulation.

Finally, to-date, the most relevant information about neuron firing sequences has been obtained with auditory stimuli (Luczak et al., 2007; Luczak et al., 2009, 2013), with a limited extension to somatosensory stimuli. To augment the generality of these findings, an extension of investigations to other modalities such as the visual domain has been warranted.

Motivated by these considerations, here we investigated cortical sequential firing patterns at different levels of anesthesia in rats *in vivo* using the inhalational anesthetic desflurane. At each anesthetic concentration we compared firing rate and latency patterns between spontaneous bursts (UP) and evoked stimulus responses (Stim). We specifically asked if brief visual stimuli elicit reproducible firing latency sequences of visual cortex neurons and if so, how do the sequences compare when stimuli are applied during DOWN vs. UP states, how they compare to those during spontaneous firing (UP state). We then asked if anesthesia at graded levels would alter these sequences and if it did, whether there were specific types of neurons, i.e., excitatory, or inhibitory, or those with low or high firing rate, contributing to the observed effects. We hypothesized that spontaneous UP and Stim sequences may be partially different and that a significant change in sequences would occur at an anesthetic dose associated with suppression of consciousness. As we will show, the anesthetic desflurane does indeed exert a dose-dependent effect on spike latencies both in UP state and post-stimulus although, contrary to our expectations, mainly at a shallow level of sedation, suggesting an early disruption of sensory information processing before behavioral signs of unconsciousness are expressed.

## MATERIALS AND METHODS

### Experimental Design

This study used data presented in a previous publication (Lee et al. 2020). Briefly, extracellular electrophysiological signals were recorded from layers 5-6 of the primary visual cortex of adult Long-Evans rats using chronically implanted 64-channel microelectrode arrays 1~8 days after implant surgery. The volatile anesthetic desflurane was administered in stepwise decreasing inhaled concentrations of 6%, 4%, 2%, and 0%, from here on indicated as C6, C4, C2 and C0, respectively. A 15-minute equilibration period was allowed at each step. Body temperature was controlled at 37°C by subfloor heating. In each condition, electrophysiological signals were recorded for 20 minutes in resting state followed by 6-minute recording with visual stimulation using discrete light flashes delivered by transcranial illumination as previously described (Galambos et al., 2001; Todorov et al., 2016). Flashes were delivered at randomized interstimulus intervals (2-4 seconds; 10ms pulse, 100 flashes per anesthetic level). In one rat, resting state recording was limited to 10 minutes. In six rats, additional flashes of 1ms pulse duration were presented but not analyzed here.

### Recording and unit classification

Signals were sampled at 30 kHz (SmartBox; Neuronexus Technologies, Ann Arbor, MI) and median referenced. To mitigate motion artifacts, data segments with absolute values greater than 10 standard deviations were removed. By this criteria, one rat was completely excluded due to excessive noise. Single unit spiking activity was determined from the 300-7500 Hz frequency band using template-based spike sorting software SpyKing Circus (Yger et al., 2018) followed by manual curation. Neurons were classified as putative inhibitory if their spike waveform had short half-amplitude width and short trough-to-peak time (Csicsvari et al., 1998). The remaining neurons were classified as putative excitatory. Spike times were down sampled to 1KHz.

### Detection of sequences

To detect and compare spiking sequences of varying biophysiological and temporal scales we needed to use distinct detection methods for C0 spontaneous theta (Theta), C2-C6 UP-states (UP), and C0-C6 visual stimulus induced responses (Stim). Using distinct detection methods may cause biases, so we applied appropriate normalization as explained later. We confirmed our conclusions by observing the raw spike raster alongside testing for statistical significance.

In the awake condition, spiking activity appears nearly continuous, and the onset of embedded bursts is difficult to detect. To overcome this difficulty, the timing of spike bursts was linked to the peaks of population firing. Spike times were summed across all neurons and convolved with a 100ms gaussian kernel. To further improve detection, only pronounced peaks were included with minimum prominence 7-8, where prominence is the peak’s height relative to other local peaks (MATLAB2021a findpeaks command). From the Theta onset, we extracted 200ms segments (100ms before and 100ms after peak) as a trial for the awake Theta spike sequence. Upon observing the detected onsets, we found falsely detected peaks occurring away from the Theta bursts. We removed these by excluding peaks not occurring in succession of each other, specifically by excluding maximal peaks not within 200ms of each other. For a similar reason, we also excluded detections with less than 4 peaks of within 1500ms. The minimum prominence of the maximal peaks was adjusted in each rat by manually checking the alignment of detected onsets and Theta bursts.

To detect the onset of spontaneous C2-C6 UP-state spike sequences, we sought to identify the moment when the UP-state began bursting spikes. We began by isolating the slow component in the population firing which includes the UP-states by band-pass filtering at 0.5-4Hz. Since band-passing centers the data to zero, we took zero-crossing from negative to positive as the candidate onsets. From these candidate onsets we determined which were associated with a large change in activity, i.e., a transition from a relative silence to UP-state. For further analysis we only included onsets with a large activity distance between minima pre-onset to maxima post-onset that were greater than 60~90th percentile. This percentile cutoff was adjusted for each rat by manually checking the onset alignment to the transition from silence to UP-state. 200ms segments were extracted from the C2-C6 UP-state onsets for analysis. For C0-C6 Stim, we segmented 200ms from visual stimulus.

For the sequence detections mentioned above we used a consistent segment size of 200ms to avoid bias by the varying durations of different sequence types. We also excluded brief periods of paradoxical desynchronized activity that occasionally occur during anesthesia during both evoked and spontaneous bursts. (Lee et al., 2020a).

### PETH and latency estimate

After extracting the segments for each sequence, we estimated each neuron’s latency within the sequence by the central mass of peri-event time histograms (PETHs) (Luczak et al., 2009). PETH was calculated using 4ms Gaussian filter followed by min-max scaling to the range of 0 to 1 using the formula:

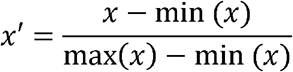

where x’ is the rescaled distribution and x is the original distribution. A relatively narrow 4ms Gaussian filter was used to emphasize the subtle dispersed spikes. For comparison among conditions, we z-scored the estimated latency to avoid biases among rats or sequences. We visualized the similarity of latency estimates between sequences (C0 Theta, C0-C6 Stim, C2-C6 UP), using multi-dimensional scaling plots which apply principal components analysis on the similarity matrix. Because 7-8Hz theta local field potential during wakefulness state have a similar period to that of the 50-200 ms spike sequences or “information packets” (Luczak, 2015), the theta cycle was used as a reference for sequences in C0. For visualizing latency estimates, the similarity matrix is the ρ^2^ value after Spearman correlation.

### Reliability

The reliability of the neuron’s latency estimate was by the similarity (correlation) between spike train trials (Schreiber et al., 2003; Luczak et al., 2007) as

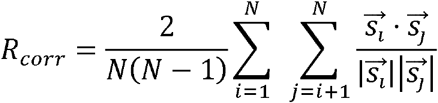

where N is the number of trials, and 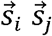 are 12ms Gaussian filtered spike train. In other words, we are correlating 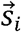 and 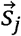 across trials i,j = 1,…,N and scaling by 2/N(N-1). The 12ms Gaussian filter is a common window size when calculating reliability (Schreiber et al., 2003; Luczak et al., 2007).

### Normalized firing rate

The firing rate of sequences was calculated by counting the number of spikes within the sequence segment and dividing by the segment length, 200ms. When comparing between sequence types and between rats, we used min-max scaled the firing rate to the range of 0 to 1. This rescaling was applied per trial. We visualized the normalized firing rates among sequence trials using multi-dimensional scaling on the distance matrix, or the trial dissimilarity, calculated by Euclidean distance applied on trial firing rates. Specifically, we used classical multi-dimensional scaling which is equivalent to applying principal component analysis to the distance matrix (MATLAB2021a cmdscale command). After applying multi-dimensional scaling, the dissimilarity of normalized firing rate among sequence trials is represented as two-dimensional distance, we call this firing rate distance.

### Statistical Analysis

When testing for variable dependence we used Spearman correlation. For testing differences between paired variables, we used Wilcoxon signed-rank test and for unpaired variables, we used Wilcoxon rank-sum test. When testing effects for two or more groups we used Friedman test, or Kruskal-Wallis one-way analysis of variance for groups with missing values. When multiple tests were used to support a conclusion, we used false discovery rate correction by Benjamini-Hochberg Procedure (Hochberg and Benjamini, 1990).

## RESULTS

### Stimulation on DOWN state evokes response with high reliability

Desflurane anesthesia at moderate to deep levels C4 – C6 produced spontaneous neuronal activity consisting of UP and DOWN states. Because flash stimuli were applied at random intervals with no respect to ongoing activity, we first asked how the pre-stimulus state, UP and DOWN, affected the stimulus-evoked spike sequences (Stim-on-DOWN, Stim-on-UP). Neurons were sorted by their estimated latency averaged between Stim-on-DOWN and Stim-on-UP from C6 data. Figure 1A shows a spike raster of sorted neurons from one experiment as an example. With all data combined, Figure 1B shows that Stim-on-DOWN PETH had well-aligned latencies, while those in Stim-on-UP PETH were more dispersed between 0 and 100ms. The estimated latencies between Stim-on-DOWN and Stim-on-UP were only weakly correlated (Figure 1C, Spearman correlation r^2^ = 0.032, p = 0.015, N = 188). Further 80% of neurons latencies of Stim-on-DOWN were delayed compared to Stim-on-UP.

**Figure 1.**
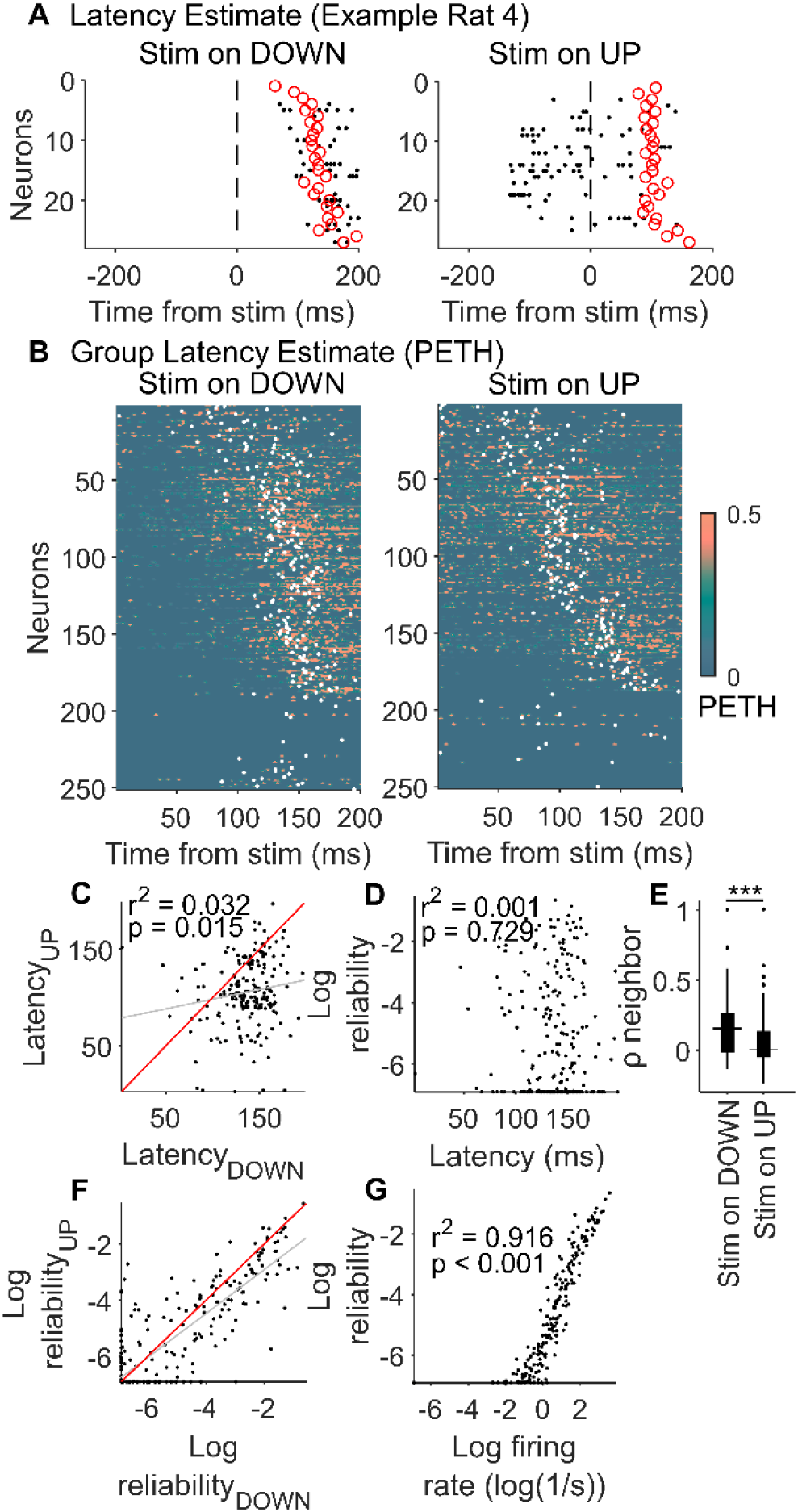
Stim on DOWN evokes response with high reliability. Evoked sequential responses after Stim on cortical DOWN and UP states are compared in anesthesia C6. A) Example of spike raster for Stim on DOWN and UP from one experiment. Red circles indicate latency estimated from the central mass of PETH. B) Group PETH distribution for Stim on DOWN and UP. PETH distribution were calculated for seven rats, then sorted by their latency estimates (white dots) averaged between Stim on DOWN and UP (trial sizes: N_DOWN_ = 73, 56, 13, 16, 39, 75, 41; N_UP_ = 17, 29, 52, 46, 41, 11 33). C) Spearman correlation between estimated latencies of Stim on DOWN and UP (neuron samples, N = 188). The red line is Latency_DOWN_ = Latency_UP_. D) Reliability versus latency (Spearman correlation, neuron samples, N = 213). E) Comparing ρ-values after correlating neighboring neuron PETHs sorted by the estimated latency (Wilcoxon rank-sum, neuron samples, N_DOWN_ = 212; N_UP_ = 202), ***p<0.001. F) Log reliability between Stim on DOWN and UP (neuron samples, N = 251). The red line is Log reliability_DOWN_ = Log reliability_UP_. G) Reliability versus firing rate (Spearman correlation, neuron samples, N = 251).

### Deep anesthesia preferentially reduces inhibitory firing

Next, we delineated how an incremental change in anesthetic level modulates the firing rate of classified excitatory and inhibitory neurons. Figure 2A illustrates by multi-dimensional scaling the gradual transformation of the firing rate of all neurons from C0 to C6 in one experiment as an example. In the awake spontaneous condition, which shows no easily discernible UP state, the theta cycle was used as a reference (C0 theta, see Methods). Further, we compared the mean Euclidean distance between firing rate trials or firing rate distance (Figure 2B). Group analysis confirmed that the firing rate distance (dissimilarity) to C6 sequences gradually increased from C6 to C0 (Figure 2C, Friedman test after averaging, x^2^ = 9.86, p = 0.0198, N = 7).

**Figure 2.**
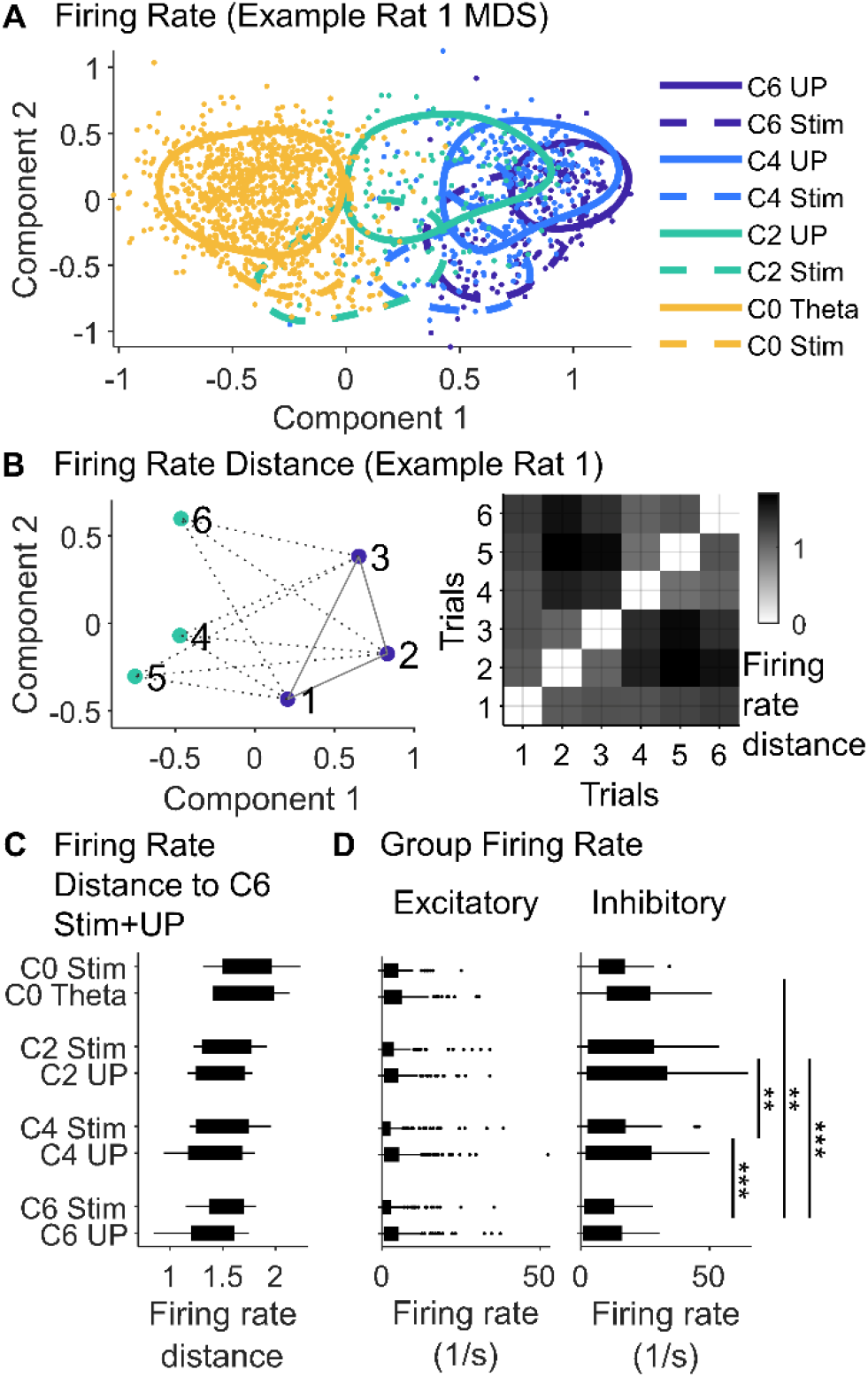
Cortical saturation of desflurane reduces inhibitory firing. Firing rate comparison across C0 to C6 desflurane states and spontaneous (UP and theta) and evoked Stim bursts. A) Multi-dimensional scale (MDS) plot of per trial normalized firing rate in one experiment as an example (N = 1571 trials). B) Left plot: example of 6 trials from above MDS visualization of firing rate distance, with grey lines representing pairwise firing rate distance between trials of C6 UP to C2 UP (dotted line) and within C6 UP (solid line). Right plot: we compare the mean Euclidean distances between trial firing rates in C. C) Firing rate distance to C6 sequences (rat samples, N = 7). D) Firing rate of sequences for excitatory and inhibitory neurons (neuron samples, N_ex_ = 216, N_in_ = 35). Wilcoxon signed-rank applied on the within state averages, false discovery rate corrected on the state comparisons, *p<0.05; **p<0.01; ***p<0.001.

When two classes of neurons were analyzed separately, we found that the anesthetic depth had no effect on excitatory cell firing (Figure 2D left, Friedman test, x^2^ = 5.97, p = 0.1129, N = 216), but suppressed inhibitory cell firing (Figure 2C right, Friedman test, x^2^ = 38.93, p < 0.001, N = 35).

### UP and Stim spike sequences change with anesthetic depth

We then delineated how the level of anesthesia affects spike latency sequences. Multi-dimensional scale plots created from data from all rats combined revealed that both spontaneous sequences (C0 Theta, C2 to C6 UP) and Stim sequences (C0 to C6 Stim) changed gradually through the anesthetic levels (Figure 3A). Repeating the latency analysis in each rat separately we found that the latencies gradually changed for both Stim (Figure 3B left, Kruskal-Wallis test, x^2^ = 12.76, p = 0.0017, N_C0,C2,C4_ = 5,7,7) and UP (Figure 3B right, Kruskal-Wallis test, x^2^ = 6.02, p = 0.0493, N_C0,C2,C4_ = 7,7,7). Followed up with Wilcoxon tests with Benjamini correction, the UP/theta state sequences were all significant but the Stim sequences were non-significant (Figure 3B).

**Figure 3.**
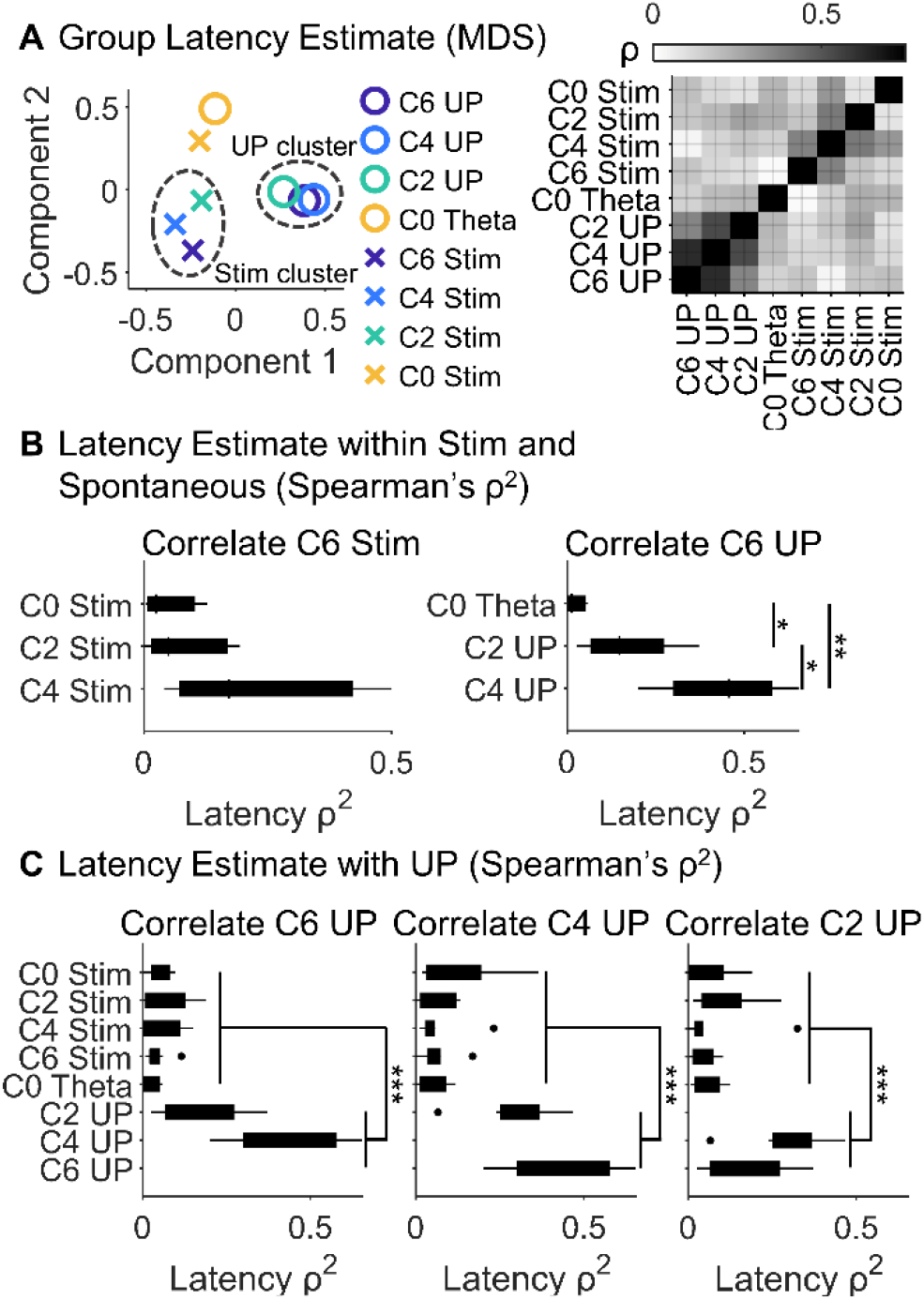
UP and Stim have distinct sequences. Latency comparison of spontaneous UP, Theta, and evoked Stim bursts across C0 to C6 states. A) Left: multi-dimensional scaling plot of Spearman correlation ρ-values between sequence latencies. UP (C2, C4, C6 UP) and Stim clusters (C2, C4, C6 Stim) are observed. Data are from seven rats. Right: distance matrix of the corresponding ρ-values. B) Spearman correlation ρ ^2^ values correlating within Stim (left, N_C0,C2,C4_ = 5,7,7) and spontaneous sequences (right, N_C0,C2,C4_ = 7,7,7). C) Spearman correlation ρ ^2^ values of latency sequences in C6, C4, C2 UP vs. the other sequences (N = 7 rats). Wilcoxon rank-sum compares the compiled UP (N = 14) versus non-UP (N = 33) bursts, false discovery rate corrected amid C6 to C2 UP, *p<0.05; **p<0.01; ***p<0.001.

Because firing rate of inhibitory neurons decreased during anesthesia, we considered the possibility that the observed latency changes were caused by those of inhibitory cell firing. To address this possibility, we repeated the latency analysis for excitatory neurons only and again found that the UP/theta sequences were significant (Kruskal-Wallis test, x^2^ = 12.03, p = 0.0024, N_C0,C2,C4_ = 5,7,7), whereas the Stim sequences were not (Kruskal-Wallis test, x^2^ = 4.21, p = 0.1220, N_C0,C2,C4_ = 7,7,7). These results support that the latency changes of spontaneous sequences were not due to the altered inhibitory firing rate.

The results from multi-dimensional scaling of the Spearman correlation ρ-values revealed two distinct clusters containing data from the anesthetized UP and anesthetized Stim sequences (Figure 3A). In addition, the awake C0 Theta and C0 Stim were distinct from both Stim and UP C2-C6 clusters. Consistently, ρ^2^ values correlating within C2/C4/C6 UP were higher than other sequences (Figure 3C, Wilcoxon rank-sum test of C2/C4/C6 UP sequence vs. other sequences, p < 0.001, N_UP_ = 14, N_other_ = 33). This suggests that the spike latency correlations in the awake condition were different from both spontaneous and stimulated spike latencies at all levels of anesthesia.

### Low firing excitatory neurons differentiate UP and Stim sequences

To understand why the UP and Stim clusters are distinct, we partitioned neurons based on their cluster participation as UP&Stim, UP&¬Stim, ¬UP&Stim, where the symbol ¬ indicates “not” (Figure 4A). Participating neurons were defined as those within 1 SD orthogonal distance from the prediction line fitted with orthogonal least squares. For example, UP&¬Stim neurons are participating in the UP cluster, and therefore within 1 SD orthogonal distance from the prediction line for C2/C4/C6 UP vs. C4/C6/C2 UP, but not the Stim cluster. We verified that neurons in each in the participation group were relatively evenly represented (rat proportion, N = 7, UP&Stim 11:19:12:18:19:16%, UP&¬Stim 10:6:9:14:30:19%, ¬UP&Stim 0:21:0:14:21:21%).

**Fig 4.**
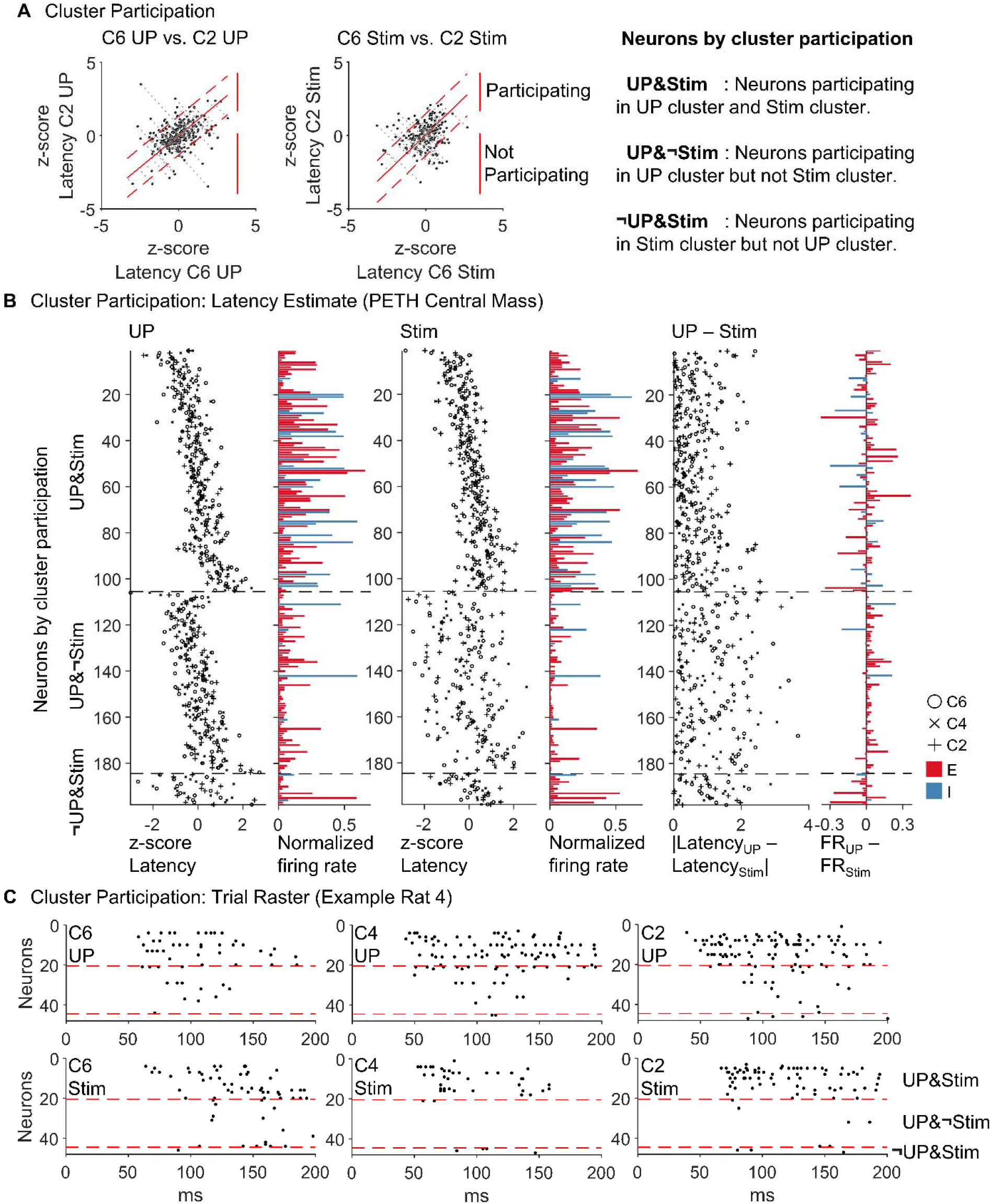
Low firing excitatory neurons differentiate UP and Stim sequences. UP (C2, C4, C6 UP) and Stim clusters (C2, C4, C6 Stim) as delineated in the multi-dimensional scaling plot of Spearman ρ^2^ between PETH central mass estimated latencies. Data are from seven rats. A) Scatter plot of neuron participation from which Spearman ρ^2^ is calculated for C2 vs C6 as an example. Neurons participate in both C2 UP and C6 UP if they are within 1 SD orthogonal distance (red dotted line) from the line of fit (red solid line). Legend on the right defines three categories of neurons: UP&Stim (N = 105), UP&¬Stim (N = 79), ¬UP&Stim (N = 14). B) Latency estimates and firing rates for UP, Stim and UP - Stim. Neurons are sorted by the three categories of cluster participation. Firing rates are normalized to min-max scale range 0~1 for each UP or Stim trial. Plotted firing rates are averages of C2, C4, C6. C) Example trial raster from rat 4. Neurons are sorted by the three categories of cluster participation.

We found that the UP&¬Stim neurons had higher variability of C2/C4/C6 Stim latencies than C2/C4/C6 UP latencies (Figure 4B UP&¬Stim, Var_UP_=0.64, Var_Stim_=1.42). Further, the UP&¬Stim group contained mostly excitatory neurons and their UP cluster had higher firing rate than the Stim cluster (Figure 4B middle, UP – Stim). Therefore, the higher reliability of UP cluster latency may be attributed to the higher firing rate of excitatory neurons, which agrees with our observation that the firing rate affects reliability (Figure 1G). In other words, firing rate is coupled not only to the reliability among sequence trials (Figure 1G), but also to the sequence consistency across anesthetic depths C2/C4/C6 (Figure 4B). The higher reliability and firing rate of ¬UP&Stim in the C2/C4/C6 Stim cluster compared to that in the C2/C4/C6 UP cluster supports this notion. Upon examining the spike rasters, we see that UP&¬Stim excitatory neurons extend the Stim sequences to form the UP sequences (Figure 4C). Conversely, the UP&Stim neurons comprise high-firing neurons with similar patterns of latency estimates between UP and Stim (Figure 4C top, UP and Stim) with lower variance in the latency difference (Figure 4B top, UP – Stim). Compared to the other categories of cluster participation, UP&¬Stim are lower firing rate and contained mostly excitatory neurons (Figure 4B). Therefore, UP and Stim sequences are differentiated by activation of low firing excitatory neurons.

## DISCUSSION

In this work we aimed to determine how anesthesia at various levels may modulate spike latency sequences during spontaneous UP states and following sensory stimulation in rodent visual cortex. It has been shown that in both conditions, neurons of a local population engage in firing with a stereotypic sequence of latencies that are conserved across trials, sensory stimuli, and cortical states (Luczak et al., 2009; Mackevicius et al., 2019; Filipchuk et al., 2022). Such stereotypic sequences have been proposed to form the basic building blocks of cortical computations underlying a range of functions from sensory feature coding to cognition (Luczak et al., 2015). Former investigations focused on the auditory cortex and auditory stimuli and emphasized the similarity of spontaneous and stimulus evoked sequences and their invariance with cortical state contrasting wakefulness and anesthesia achieved mostly with urethane. Corresponding investigations in the rodent visual system and using a clinically used inhalational anesthetic of different pharmacology titrated to graded anesthetic levels have not been performed.

By recording extracellular single unit activity in the rat visual cortex *in vivo*, we analyzed the spiking structure of spontaneous UP bursts (UP) and visual flash-evoked responses (Stim) during stepwise changes of the inhaled concentration of the anesthetic desflurane. We reached three main conclusions. 1) Stimuli applied during silent periods (DOWN state) produce cortical spike responses with high reliability and increased delay compared to spike responses after UP. 2) Anesthesia alters UP and Stim spike latency sequences by suppressing the firing of inhibitory neurons. 3) Latency patterns occurring spontaneously and by stimulation are similar in wakefulness, but become progressively different in anesthesia, and 4) Overall firing rate is higher in UP than in Stim due to the recruitment of excitatory neurons.

Former studies found that flashed bar stimuli presented during electrical silence evoke larger magnitude of intracellular voltage (Haider et al., 2013). Another study also found whisker deflection during DOWN state evoked large and delayed responses compared to stimuli during the UP state (Petersen et al., 2003). Our finding confirms these showing that spike response patterns were more robust and reliable when stimuli were applied during DOWN vs. UP periods. This difference may in part be due to that stimuli presented during UP states vs Down states elicit responses that are temporally and spatially more restricted (Petersen et al., 2003). It was suggested that UP/DOWN states alter the evoked responses by primarily altering internal neuronal state, rather than network level interactions (Hasenstaub et al., 2007). Our results support the network level interactions, as the UP state disrupts the response sequence by lowering reliability and stimuli on the DOWN state evoked responses which modulates the sequence by stimuli and anesthetic infusions.

We also found that desflurane anesthesia exerted a differential effect on visual cortex neurons, suppressing the firing of putative inhibitory neurons but not excitatory neurons. Anesthetics diminish the efficiency of feed-forward inhibitory circuits (Gabernet et al., 2005), shrink the receptive field size of thalamic relays (Katz et al., 2012), and reduce cortical IPSPs by 90% while sustaining cortical EPSPs at 50% (Taub et al., 2013). Similar to the Stim sequences, UP sequences showed decreased inhibitory firing rate with anesthetic depth. Previous studies showed that spontaneous UP states can be initiated from both thalamic and cortical sources suggesting the potential involvement of multiple mechanisms (Murphy et al., 2009). Along with suppressing thalamocortical input, anesthetics could also influence UP-state sequences through intracortical synaptic and cellular changes, such as the postsynaptic facilitation or inhibition, modification of two-pore potassium channels and NMDA receptors (Alkire et al., 2008), as well as ATP-dependent cellular energy depletion (Ching et al., 2012).

Prior studies also found that the Stim firing rate distribution was nested within the UP firing rate distribution (Luczak et al., 2022). Our results did not reproduce this finding. This discrepancy could be due to the difference in preprocessing methods as Luczak et al. included in the analysis only neurons with firing rate higher than 2 Hz. In contrast, our analysis attributed these low firing excitatory neurons to differentiate UP and Stim sequences, which may further the discrepancy. Recent studies using two-photo calcium imaging with estimated raster also exclude cortical low firing neurons and found anesthesia spontaneous and stimuli evoked sequence were overlapped (Filipchuk et al., 2022), supporting the interpretation that spontaneous and evoked sequence overlap is related to the preprocessing of low firing rate neurons. On one hand, this preprocessing method has resulted in finding sequence mechanisms specific to high firing neurons, which for example selectively respond to both simple and high order stimuli (Luczak et al., 2022) or altering state to behavior (Okun et al., 2015). Excluding low firing rate neurons also tend to reduce dimensionality and enhance the statistical power of the analysis (Altman and Krzywinski, 2018). The low firing neurons also reveal unique functions with precise and selective response to sensory stimuli (Bachatene et al., 2015; Clawson et al., 2018; Lee et al., 2020b). Both the exclusion and inclusion of low firing rate neurons, therefore, reveal unique aspects of the cortical spiking activity. The discrepancy may also be due urethane and desflurane having markedly different pharmacological profiles, due to a difference in both anesthetic type and dose. Desflurane alters the excitatory-inhibitory balance of both spontaneous UP and evoked Stim sequential bursts by augmenting the excitatory response in the background of increased inhibitory tone (Lukatch et al., 2005; Hudetz and Imas, 2007; Vizuete et al., 2012).

Lastly, we delineated how the anesthesia at various levels influenced the spike sequence latency in UP and Stim. For the awake condition C0, the theta cycle was used as an endogenous temporal reference. It was previously noted that during active waking, 7-8Hz theta of the local field potential dominates that has a similar period to that of the 50-200 ms spike sequences or “information packets” (Luczak et al., 2015). Place cells in the rodent hippocampus also fire sequences aligned with theta cycle (Diba and Buzsaki, 2007). Spontaneous sequence C0 Theta and C0 Stim had similar latency patterns but dissociated as the anesthesia was made deeper. In anesthesia, spike pattern in UP was different from that in Stim by added spike from low firing excitatory neurons. These results were reproduced for excitatory neurons alone, demonstrating the latency findings are independent of the anesthetic induced suppression of inhibitory firing rate. The similarity in latency patterns between C0 Theta and C0 Stim suggests that integration of both spontaneous and stimulus-related information was effective in the awake state. This contrasted with anesthesia, which presumably disconnects the stimulus response from ongoing cortical activity, as implied by the observed difference between spontaneous and evoked latency patterns. Of note is that for both UP and Stim, a distinct change in latency clusters occurred between C0 and C2 with minimal further change from C2 to C6. In C2, the animals were mildly sedated but behaviorally responsive and presumably conscious, whereas in C6 they were unresponsive and arguably unconscious. Thus, the change in spike patterns from C0 to C2 was likely associated with a drop in the animals’ level of vigilance rather than their level of consciousness that normally commences around C4 as indicated by the appearance of slow electrocortical oscillations (Alkire et al., 2008) and the loss of the animals’ righting reflex (Imas et al., 2005b; Vizuete et al., 2012).

The distinct sequential patterns in UP vs. Stim at all examined anesthetic levels, was attributed to low-firing excitatory neurons contributing to the UP sequence. Previous studies suggested that sparsely firing excitatory neurons, rather than high-firing neurons, selectively respond to a variety of stimuli (Clawson et al., 2018). In Sweeney and Clopath’s mathematical model, low-firing, low-population coupling neurons function as a backbone stimulus representation, driven by slow synaptic change, while high-firing, high-population coupling neurons function as a flexible substrate driven by fast synaptic changes (Bachatene et al., 2015; Sweeney and Clopath, 2020). Furthermore in our previous study, the role of low/high firing and low/high population coupling was sustained in graded anesthetic states (Lee et al., 2020b), suggesting the function of neurons remained intact in anesthesia. Our results and previous studies suggest low-firing excitatory neurons have unique role to learning by synaptic modification in neuron networks (Bachatene et al., 2015; Sweeney and Clopath, 2020).

It has been proposed that the organized firing of neurons during the first 50-200ms after stimulus or onset of spontaneous Up state represents a basic building block of cortical coding and communication (Luczak et al., 2009). Specifically, the early response within 100ms, which is more stereotypic, is thought to encode simple stimulus properties, whereas the later response, which is more variable, encodes complex features (Luczak et al., 2015). The late response may be governed in part by long range circuit interactions including top-down feedback. Interestingly, anesthetics preferentially suppress the late response of visual cortex neurons specifically at a drug concentration that leads to loss of consciousness (Hudetz et al., 2009) and this is paralleled by a reduction of top-down feedback (Imas et al., 2005a) consistent with the proposal that the late parts of neuronal coding may be essential to sustaining conscious sensory perception (Mashour et al., 2020). The modification of spike sequences during anesthesia as observed here represents another, finer signature of altered sensory processing in visual cortex.

## ACKNOWLEDGEMENTS

Research reported in this publication was supported by the National Institute of General Medical Sciences of the National Institutes of Health under award number R01-GM056398 and the Center for Consciousness Science, Department of Anesthesiology, University of Michigan Medical School, Ann Arbor, Michigan, USA. The content is solely the responsibility of the authors and does not necessarily represent the official views of the National Institutes of Health. The authors thank Kathy Zelenock, MS for her assistance in laboratory operations.

